# Synthesis of perfluorooctanoic acid-containing membrane lipids by human pathobionts

**DOI:** 10.1101/2025.10.14.682380

**Authors:** Ziqiang Guan, Aparna Uppuluri, Guan Chen, Kelli L. Palmer

## Abstract

Per- and polyfluoroalkyl substances (PFAS) are synthetic fluorinated compounds used widely in industrial and consumer products. They are unusually stable due to carbon–fluorine bonds and resistant to degradation, making them persistent contaminants in water, soil, and biota. PFAS are associated with adverse health effects in humans including cancer and liver disease. The effects of PFAS on human-associated bacteria are largely unexplored, a significant gap in knowledge because these bacteria are exposed to PFAS *in vivo* at sites including the colon and bladder. One of the best studied PFAS compounds is perfluorooctanoic acid (PFOA), an eight-carbon perfluorinated carboxylic acid whose structure is analogous to a fatty acid. Here, we cultured *Enterococcus faecalis*, a Gram-positive bacterium, and *Pseudomonas aeruginosa*, a Gram-negative bacterium, in growth medium supplemented with PFOA and corresponding control conditions and performed lipidomic analyses using liquid chromatography-tandem mass spectrometry (LC-MS/MS) to elucidate lipid remodeling in response to PFOA exposure. Strikingly, novel fluoroalkyl-containing membrane lipids are synthesized by both of these bacteria, with each species synthesizing unique fluoroalkyl-lipids. Moreover, a high-level daptomycin-resistant strain of *E. faecalis* produces strikingly high levels of fluoroalkyl-lipids, demonstrating that prior antibiotic exposure and concomitant effects on bacterial evolution can alter bacterial interactions with PFAS. Because bacterial lipids are important immunomodulators *in vivo*, we propose that PFAS-containing bacterial lipids may be novel mediators of host–microbe–pollutant interactions. Our results also establish a novel mechanism for the bioaccumulation of PFOA and, potentially, for bioremediation of PFOA in biological systems such as the human gastrointestinal tract.

## Importance

PFAS are widespread pollutants whose effects on human health are a very active area of investigation. Bacteria colonize humans and can profoundly impact human health. Bacteria are exposed to PFAS in the human body, but the effects of PFAS on bacteria are poorly understood. Here, we used comprehensive analysis of bacterial lipids to study whether PFAS exposure alters lipid biosynthesis and chemistry in bacteria, using two human pathogens as models for our study. We found that both bacterial species appear to scavenge PFAS from their environment and use them as building blocks for lipid synthesis. This is a significant finding because it suggests that human-associated bacteria may alter human interactions with PFAS via transformation of PFAS and via synthesis of PFAS-containing metabolites. These novel lipids are likely to impact host-pathogen interactions and bacterial evolution, which should be investigated in future studies.

### Manuscript body

#### Per- and poly-fluoroalkyl substances (PFAS) are persistent pollutants

PFAS are widely used in commercial products and deemed emerging persistent pollutants by the Stockholm Convention(1). The extraordinary stability of their carbon-fluorine bonds makes PFAS recalcitrant to degradation(2), thus PFAS can bioaccumulate in food chains(3). The effects of PFAS on biological systems are multifactorial and remain poorly understood for the thousands of PFAS structural variants available(2, 3). PFAS have been linked to adverse health effects including cancers and liver disease(3). Of relevance for our study, long-chain PFAS (e.g., perfluorooctanoic acid [PFOA], **Fig. 1A**) possess physiochemical properties of aliphatic chain compounds which interact with lipid membranes(4).

**Figure 1.**
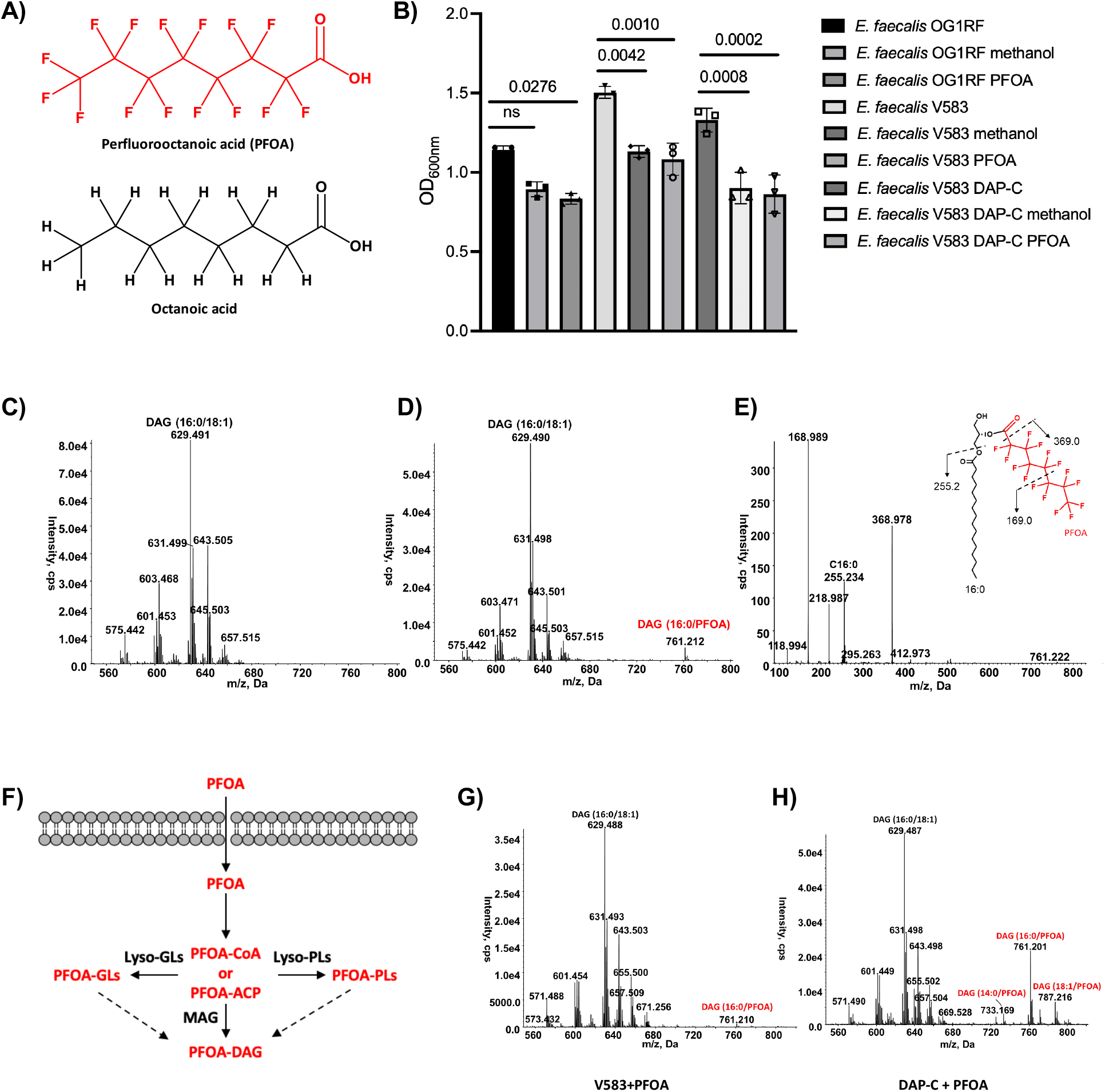
Detection of PFOA-containing diacylglycerol (DAG) in *E. faecalis*. A) Chemical structures of PFOA and octanoic acid. B) Effects of PFOA on the culture growth of *E. faecalis*. Bars on graph represent the average overnight growth yield (as measured by optical density at 600 nm (OD600nm) across independent replicates (n=3). The bars represent mean with standard eviation. Significance was assessed using ordinary one-way ANOVA for multiple comparisons. Negative ion ESI mass spectrum showing [M+Cl]^−^ ions of DAG molecular species in *E. faecalis*. Synthesis of DAG (16:0/PFOA) in *E. faecalis* treated with PFOA. E) Structural confirmation of DAG (16:0/PFOA) by MS/MS. F) Proposed mechanisms of PFOA-DAG formation in *E. faecalis*. The process involves initial transport of PFOA across the cell membrane. PFOA is then converted into metabolically active forms, PFOA-CoA or PFOA-ACP, which serve as substrates for acyltransferases to synthesize PFOA-containing membrane lipids. These lipids can be further metabolized to form PFOA-DAG. It is unclear why PFOA is specifically enriched in DAG. G) Low-level PFOA-containing DAG is detected in *E. faecalis* V583 treated with PFOA. H) High-level PFOA-containing DAG species are detected in PFOA-treated DAP-C, a daptomycin-resistant strain evolved from V583.

#### Little is known about response of bacterial pathogens to PFAS

Humans encounter PFAS via routes including diet and lung inhalation; PFAS are detected in human tissues and body fluids; and PFAS are excreted via the bladder or colon(3, 5). Thus, bacteria that colonize these sites are exposed to PFAS *in vivo*. The ramifications for these exposures on bacterial physiology and evolution are largely unknown. In the most comprehensive study, Lindell, et al, surveyed responses of human intestinal commensal bacteria to PFAS(6). The investigators found that some species (e.g. *Bacteroides uniformis*) accumulate PFAS inside their cells while others (e.g., *Escherichia coli*) pump out PFAS via membrane export. Moreover, *B. uniformis* adapted to serial PFAS exposure, acquiring enhanced PFAS tolerance and mutations in genes with predicted roles in membrane transport and cell wall metabolism. This study provides compelling evidence that human-associated bacteria exhibit specific responses to PFAS, allowing for their survival and adaptation in the presence of these pollutants.

Here, we applied comprehensive lipidomics to investigate the response of two bacterial pathogens, *Enterococcus faecalis* and *Pseudomonas aeruginosa*, to PFOA. To our knowledge, bacterial responses to PFAS have not been evaluated by comprehensive lipidomics. This is an important gap in knowledge, given the fatty acid-like nature of PFAS, such as PFOA, which is a structural analogue of octanoic (caprylic) acid (**Fig. 1A**).

#### PFOA is incorporated into *E. faecalis* membrane lipids

Enterococci are intestinal pathobionts causing infections including endocarditis and urinary tract infections (UTI)(7). Enterococci colonize PFAS-contaminated host (gastrointestinal tract, bladder, bloodstream) and external environments (wastewater, soils), thus are ecologically-relevant model organisms(8). Remarkably, lipidomic analysis of exponentially-growing *E. faecalis* OG1RF(9) treated with 1 mg/mL PFOA for 15 minutes identified a PFOA-containing diacylglycerol (DAG) molecule at a low level (See Text S1 for Materials and Methods for our study). We then cultured *E. faecalis* strains overnight with 1 mg/mL PFOA (**Fig 1B**). A substantial level of fluoroalkyl-containing DAG was detected in these cultures by LC/MS (**Fig. 1C**). The observed [M+Cl]^−^ ion peak at *m/z* 761.212 corresponds to DAG containing C16:0 and PFOA acyl chains (predicted [M+Cl]^−^ ion at *m/z* 761.209). This peak is absent in *E. faecalis* treated with solvent only (**Fig. 1C-D**). The proposed structure of DAG(16:0/PFOA) is supported by exact mass measurement and tandem MS (MS/MS) fragmentation (**Fig. 1E**). To our knowledge, the cellular incorporation of PFAS into membrane lipids has not been previously described. Non-enzymatic acylation with PFOA is unlikely; the acylation reaction is likely catalyzed by *E. faecalis* acyltransferases (**Fig. 1F**).

#### Fluoroalkyl-containing lipids are strikingly abundant in daptomycin-resistant *E. faecalis*

The antibiotic daptomycin (DAP) is critical for treatment of enterococcal infections. DAP targets the bacterial membrane by interacting with anionic phospholipids, ultimately disrupting cellular functions(10). Enterococci can evolve DAP resistance, resulting in strains with altered lipidomes and mutations in genes associated with lipid synthesis and remodeling.(10) High-level DAP-resistant derivatives of *E. faecalis* V583(11) were previously derived, among which the strain DAP-C has 5 mutations relative to its V583 parent(12). Fluoroalkyl-containing lipids are strikingly abundant in DAP-C after culture with PFOA, including multiple variant DAGs, which are not observed for the isogenic parent V583 (**Fig. 1G-H**). These results establish that synthesis of fluoroalkyl-containing lipids by *E. faecalis* can vary based on strain genotype and prior adaptation to antibiotic exposure.

#### PFOA is incorporated into *P. aeruginosa* membrane lipids

*P. aeruginosa* causes pulmonary, wound, UTI, and other infections(13) and like *E. faecalis*, would be exposed to PFAS in a range of environments. Lipidomic analysis of *P. aeruginosa* PA14(14) treated with 1 mg/mL PFOA (**Fig. 2A**) identified the novel PFOA-containing lipid, PFOA-hydroxy fatty acid (HFA) (**Fig. 2B-D**), whose level is comparable to those of the major phospholipids phosphatidylglycerol and phosphatidylethanolamine. The synthesis of PFOA-HFA is likely related to the synthesis of hydroxyalkanoyloxy-alkanoic acid (HAA), which is the lipid moiety of rhamnolipids in *P. aeruginosa*(15) (**Fig. 2E**). This novel finding demonstrates that incorporation of PFOA into membrane lipids occurs more broadly in pathogenic bacteria.

**Figure 2.**
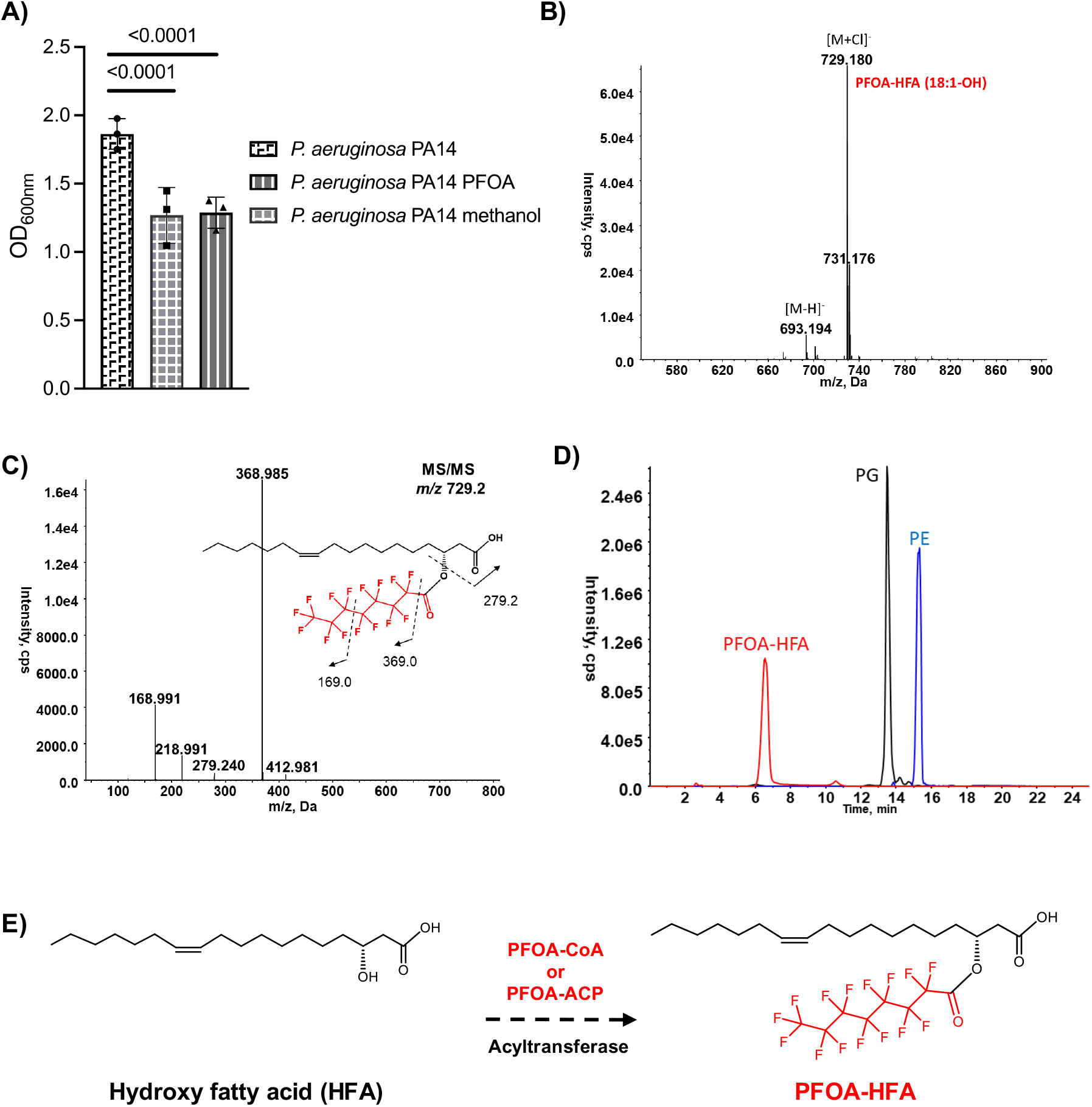
Detection of PFOA-acylated hydroxy fatty acid (PFOA-HFA) in *P. aeruginosa*. A) Effects of PFOA on the culture growth of *P. aeruginosa* PA14. Bars on graph represent the average overnight growth yield (as measured by optical density at 600 nm (OD600nm) across independent replicates (n=3). The bars represent mean with standard deviation. Significance was assessed using ordinary one-way ANOVA for multiple comparisons. B) Synthesis of PFOA-HFA in *P. aeruginosa* PA14 treated with PFOA. The deprotonated [M-H]^−^ and chloride adduct [M+Cl]^−^ ions of PFOA-HFA (18:1-OH) are detected by negative ion ESI/MS. C) MS/MS confirmation of PFOA-HFA (18:1-OH) with [M+Cl]^−^ ion at *m/z* 729.2. The double bond position depicted on the HO-C18:1 acyl chain is not determined by MS/MS and is for illustrative purposes. D) Extracted ion chromograms of major species of PFOA-HFA, PG and PE in *P. aeruginosa* PA14 treated with PFOA. E) A proposed mechanism of PFOA-HFA formation. The conversion of hydroxy fatty acid (HFA) into PFOA-HFA is catalyzed by an acyltransferase using PFOA-CoA or PFOA-ACP as a substrate.

#### Conclusions and unanswered questions

Our discovery of PFAS-containing lipids in *E. faecalis* and *P. aeruginosa* is significant and opens new research directions. Bacterial lipids, beyond lipopolysaccharide, are signaling molecules that modulate the host immune response and other processes(16). PFAS-containing bacterial lipids may be novel mediators of host–microbe– pollutant interactions *in vivo* and key missing factors for understanding the multifactorial effects of PFAS in humans, which requires investigation. Of note, different fluoroalkyl-containing lipids were detected for *E. faecalis* and *P. aeruginosa*, likely due to underlying differences in their lipid biosynthetic pathways. Future studies should analyze human-associated bacteria more broadly to catalog the diversity of fluoroalkyl-lipid molecules synthesized. The incorporation of PFAS into bacterial lipids also provides a novel mechanism for PFAS bioaccumulation and potential bioremediation *in vivo*.

Additional areas for investigation include the mechanisms of fluoroalkyl-lipid synthesis per models proposed above, and the impacts of these lipids on bacterial cell function and evolution. For example, *E. faecalis* adapts to stress via mutation(17). The replacement of the commonly occurring 18:1 side chain at the sn-2 position of DAG with the shorter side chain from PFOA is expected to have substantial effects on membrane fluidity and function. We hypothesize that, over time, the presence of fluoroalkyl-lipids in the membrane will select for compensatory adaptations that enable cells to tolerate membrane alterations caused by these lipids. These adaptations may alter antimicrobial susceptibility of *E. faecalis*.

Future studies should also explore the conditions in which bacteria synthesize fluoroalkyl-containing lipids, assessing impact of PFAS concentration, structure, and presence of host-derived fatty acids on this process. PFAS levels vary across environments; for example, PFAS in the low ng/mL range are detected in human serum for the general United States population(18) and that used here, 1 mg/mL, is comparable to the highest concentration of PFAS detected in biosolids(19). Notably, our work with DAP-C demonstrates that few mutations between otherwise isogenic strains can result in large differences in fluoroalkyl-containing lipid abundance with the same degree of PFAS exposure. Regarding PFAS structure, PFOA and other PFAS with a carboxylic acid “head group” may be misidentified as fatty acids and utilized in lipid biosynthesis by endogenous acyltransferases. If so, then PFAS of different C-F chain length and identical “head group” to PFOA may be incorporated into bacterial lipids, but PFAS with different “head groups” may not. Finally, host-derived fatty acids may outcompete PFOA as substrates if they are preferentially utilized; alternatively, their presence may be required to induce scavenging systems that inadvertently utilize PFOA.

Bacterial incorporation of PFOA into membrane lipids suggests potential parallels in mammalian cells, given shared lipid biochemistry. Comprehensive lipidomic analysis of PFOA-treated human cells should be performed to determine whether novel fluoroalkyl-containing lipids are observed.

## Acknowledgements

This work was supported by R01AI178692 and R01AI148366 to Z.G. and K.P., and the UT-Dallas Sustainability Seed Program and the Cecil and Ida Green Chair in Systems Biology Science to K.P. The Palmer lab thanks Dr. Sheel Dodani and her lab members for engaging discussions on PFAS.

## Supplementary Text S1

### Materials and methods

*E. faecalis* were cultured at 37°C in brain heart infusion medium without agitation. *P. aeruginosa* PA14^(1)^ was cultured at 37°C in Lysogeny Broth with agitation (250 rpm). For lipidomic experiments, 1 mg/mL PFOA (Millipore Sigma) in HPLC-grade methanol (Millipore-Sigma) or an equivolume amount of methanol or culture medium (each 2.7% vol/vol) was added to culture media to 15 mL final volume in polypropylene conical tubes (Thermo Fisher Scientific). 50 mL conicals were used for *P. aeruginosa* PA14 and 15 mL conicals were used for *E. faecalis*. Cultures were inoculated with single colonies and incubated for ∼18 hours. Lipid extraction was performed on 14 mL overnight culture by a modified method of Bligh and Dyer, and lipidomic analysis were performed as described previously^(2-4)^.

## Notes

### Competing Interest Statement

The authors have declared no competing interest.

## References Cited

1. Stockholm Convention on Persistent Organic Pollutants (POPs). 2024. PFASs listed under the tockholm Convention. https://chm.pops.int/Implementation/IndustrialPOPs/PFAS/Overview/tabid/5221/Default.aspx.

2. Cousins IT, DeWitt JC, Glüge J, Goldenman G, Herzke D, Lohmann R, Ng CA, Scheringer M, Wang Z. 2020. The high persistence of PFAS is sufficient for their management as a chemical class. Environmental Science: Processes & Impacts 22:2307–2312.

3. Yeoh CSL, Alrazihi LA, Wong ST, Wong SF. 2025. Per- and poly-fluoroalkyl substances (PFAS) and human health: a review of exposure routes and potential toxicities across the lifespan. Environ Toxicol Chem 44:2754–2786.

4. Jing P, Rodgers PJ, Amemiya S. 2009. High lipophilicity of perfluoroalkyl carboxylate and sulfonate: implications for their membrane permeability. J Am Chem Soc 131:2290–6.

5. Andersson AG, Fletcher T, Xu Y, Karrman A, Pineda D, Nilsson CA, Lindh CH, Jakobsson K, Li Y. 2025. The relative importance of fecal and urinary excretion of perfluorooctane sulfonic acid and perfluorooctanoic acid after high exposure - An observational study in Ronneby, Sweden. Environ Res 285:122487.

6. Lindell AE, Griesshammer A, Michaelis L, Papagiannidis D, Ochner H, Kamrad S, Guan R, Blasche S, Ventimiglia LN, Ramachandran B, Ozgur H, Zelezniak A, Beristain-Covarrubias N, Yam-Puc JC, Roux I, Barron LP, Richardson AK, Martin MG, Benes V, Morone N, Thaventhiran JED, Bharat TAM, Savitski MM, Maier L, Patil KR. 2025. Human gut bacteria bioaccumulate per- and polyfluoroalkyl substances. Nat Microbiol 10:1630– 1647.

7. Agudelo Higuita NI, Huycke MM. 2014. Enterococcal Disease, Epidemiology, and Implications for Treatment. In Gilmore MS, Clewell DB, Ike Y, Shankar N (ed), Enterococci: From Commensals to Leading Causes of Drug Resistant Infection, Boston.

8. Lebreton F, Willems RJL, Gilmore MS. 2014. Enterococcus Diversity, Origins in Nature, and Gut Colonization. In Gilmore MS, Clewell DB, Ike Y, Shankar N (ed), Enterococci: From Commensals to Leading Causes of Drug Resistant Infection, Boston.

9. Gold OG, Jordan HV, van Houte J. 1975. The prevalence of enterococci in the human mouth and their pathogenicity in animal models. Arch Oral Biol 20:473–7.

10. Nguyen AH, Hood KS, Mileykovskaya E, Miller WR, Tran TT. 2022. Bacterial cell membranes and their role in daptomycin resistance: A review. Front Mol Biosci 9:1035574.

11. Sahm DF, Kissinger J, Gilmore MS, Murray PR, Mulder R, Solliday J, Clarke B. 1989. In vitro susceptibility studies of vancomycin-resistant Enterococcus faecalis. Antimicrob Agents Chemother 33:1588–91.

12. Palmer KL, Daniel A, Hardy C, Silverman J, Gilmore MS. 2011. Genetic basis for daptomycin resistance in enterococci. Antimicrob Agents Chemother 55:3345–56.

13. Letizia M, Diggle SP, Whiteley M. 2025. Pseudomonas aeruginosa: ecology, evolution, pathogenesis and antimicrobial susceptibility. Nat Rev Microbiol doi:10.1038/s41579-025-01193-8.

14. Rahme LG, Stevens EJ, Wolfort SF, Shao J, Tompkins RG, Ausubel FM. 1995. Common virulence factors for bacterial pathogenicity in plants and animals. Science 268:1899–902.

15. Soberon-Chavez G, Gonzalez-Valdez A, Soto-Aceves MP, Cocotl-Yanez M. 2021. Rhamnolipids produced by Pseudomonas: from molecular genetics to the market. Microb Biotechnol 14:136–146.

16. Ryan E, Joyce SA, Clarke DJ. 2023. Membrane lipids from gut microbiome-associated bacteria as structural and signalling molecules. Microbiology (Reading) 169:001315.

17. Garcia-Solache M, Rice LB. 2019. The Enterococcus: a Model of Adaptability to Its Environment. Clin Microbiol Rev 32:e00058–18.

18. Sonnenberg NK, Ojewole AE, Ojewole CO, Lucky OP, Kusi J. 2023. Trends in Serum Per- and Polyfluoroalkyl Substance (PFAS) Concentrations in Teenagers and Adults, 1999-2018 NHANES. Int J Environ Res Public Health 20:6984.

19. Saliu TD, Sauvé S. 2024. A review of per- and polyfluoroalkyl substances in biosolids: geographical distribution and regulations. Frontiers in Environmental Chemistry 5:10.3389/fenvc.2024.1383185.

## References

1. Rahme, L. G., Stevens, E. J., Wolfort, S. F., Shao, J., Tompkins, R. G., and Ausubel, F. M. (1995) Common virulence factors for bacterial pathogenicity in plants and animals Science 268, 1899–1902

2. Bligh, E. G., and Dyer, W. J. (1959) A rapid method of total lipid extraction and purification Can J Biochem Physiol 37, 911–917

3. Joyce, L. R., Manzer, H. S., da, C. M. J., Villarreal, R., Nagao, P. E., Doran, K. S. et al. (2022) Identification of a novel cationic glycolipid in Streptococcus agalactiae that contributes to brain entry and meningitis PLoS Biology 20, e3001555

4. Christensen, P. M., Martin, J., Uppuluri, A., Joyce, L. R., Wei, Y., Guan, Z. et al. (2024) Lipid discovery enabled by sequence statistics and machine learning Elife 10.7554/eLife.94929.2

